# The *Saccharomyces cerevisiae* cell lysis mutant *cly8* is a temperature-sensitive allele of *ESP1*

**DOI:** 10.1101/2024.10.23.619872

**Authors:** Christopher D. Putnam, Bin-zhong Li, Lori Broderick

## Abstract

The *Saccharomyces cerevisiae* mutants designated *cly* (<u>c</u>ell <u>ly</u>sis) cause cell lysis at elevated temperatures. The *cly8* mutation, previously localized to an 80 kilobase region between *GCD2* and *SPT6* on the right arm of chromosome VII, has been used for mutation mapping and in recombination assays, but its genetic identity has remained unknown. Whole genome sequencing of *cly8* mutant and *CLY8* wild-type strains revealed four missense mutations specific to the *cly8* mutant strain in the *GCD2*-*SPT6* interval, three of these missense mutations affected essential genes. The *esp1-G543E* mutation, but not the two other missense mutations, recapitulated the temperature-sensitive growth, the cell lysis phenotype, and recessive nature of the *cly8* mutation. The *ESP1* gene encodes *S. cerevisiae* separase and is required for normal chromosome segregation during mitosis. Shifting cells containing the *esp1-G543E* mutation to the non-permissive temperature caused chromosome missegregation based on the analysis of DNA content by flow cytometry analysis. Taken together, these results indicate that the temperature-sensitive *cly8* mutation is an allele of the essential *ESP1* gene.

## INTRODUCTION

*Saccharomyces cerevisiae* mutations that result in cell lysis after prolonged incubation at 36°C have been isolated ^1-3^, and eight strains containing different mutations (*cly1-8*, called *cly* for cell lysis) were deposited with the Yeast Genetic Stock Center. Since that time, other temperature-sensitive cell lysis mutations have been isolated, including *srb1-1* and other *cly* mutations ^4-7^. Although many of these mutations remain poorly characterized, chromosomal loci have been identified for some of these mutations through genetic mapping or through complementation ^4,5,8-10^. Several genes affected by these mutations are now known: *cly4* is a mutation of *GPI2*, which encodes an enzyme involved in the synthesis of glycosylphosphatidylinositol anchors that act in cell wall structure and maintenance; *srb1-1* is a mutation of the gene *PSA1*, which is involved in cell wall biosynthesis; and *cly5, cly7*, and *cly15* are mutations of *PKC1*, which encodes protein kinase C and is essential for cell wall remodeling during growth ^4,5,9,10^.

The *cly8* mutation causes substantial growth inhibition at 30°C and complete loss of growth at temperatures above 30°C ^11^. This mutation has been localized by genetic mapping to the right arm of chromosome VII ^3,8,12-18^. The *cly8* mutation has also been used to avoid complications arising from chromosome loss in a diploid chromosome VII-based mitotic recombination assay, since loss of the chromosome containing wild-type *CLY8* gives rise to cells that cannot grow at 30°C and thus are not scored ^19-21^. Despite its utility, the gene affected by the *cly8* mutation has not been identified, which complicates targeted introduction of this mutation into new strain backgrounds. We therefore sequenced the genomes of haploid *cly8* and *CLY8* strains and a diploid *cly8/CLY8* strain generated by mating these strains, analyzed the region of chromosome VII where *cly8* has been genetically mapped, and found that the *esp1-G543E* mutation gives rise to a temperature-sensitive cell-lysis phenotype.

## RESULTS

### The *cly8* mutation is recessive and temperature sensitive

We confirmed the temperature sensitivity and recessive nature of the *cly8* mutation by spotting serial dilutions of a *cly8* haploid strain (NLBL1), a *CLY8* haploid strain (NLBL3), and a *CLY8/cly8* heterozygous diploid strain (RDKY1340) that was created by mating NLBL1 and NLBL3 at 23°C, 30°C, and 37°C ^19,21^ **(Table 1)**. As expected, only the *cly8* haploid strain showed temperature sensitivity, and this strain showed growth defects at both 30°C and 37°C **(Fig. 1)**.

**Table 1.**
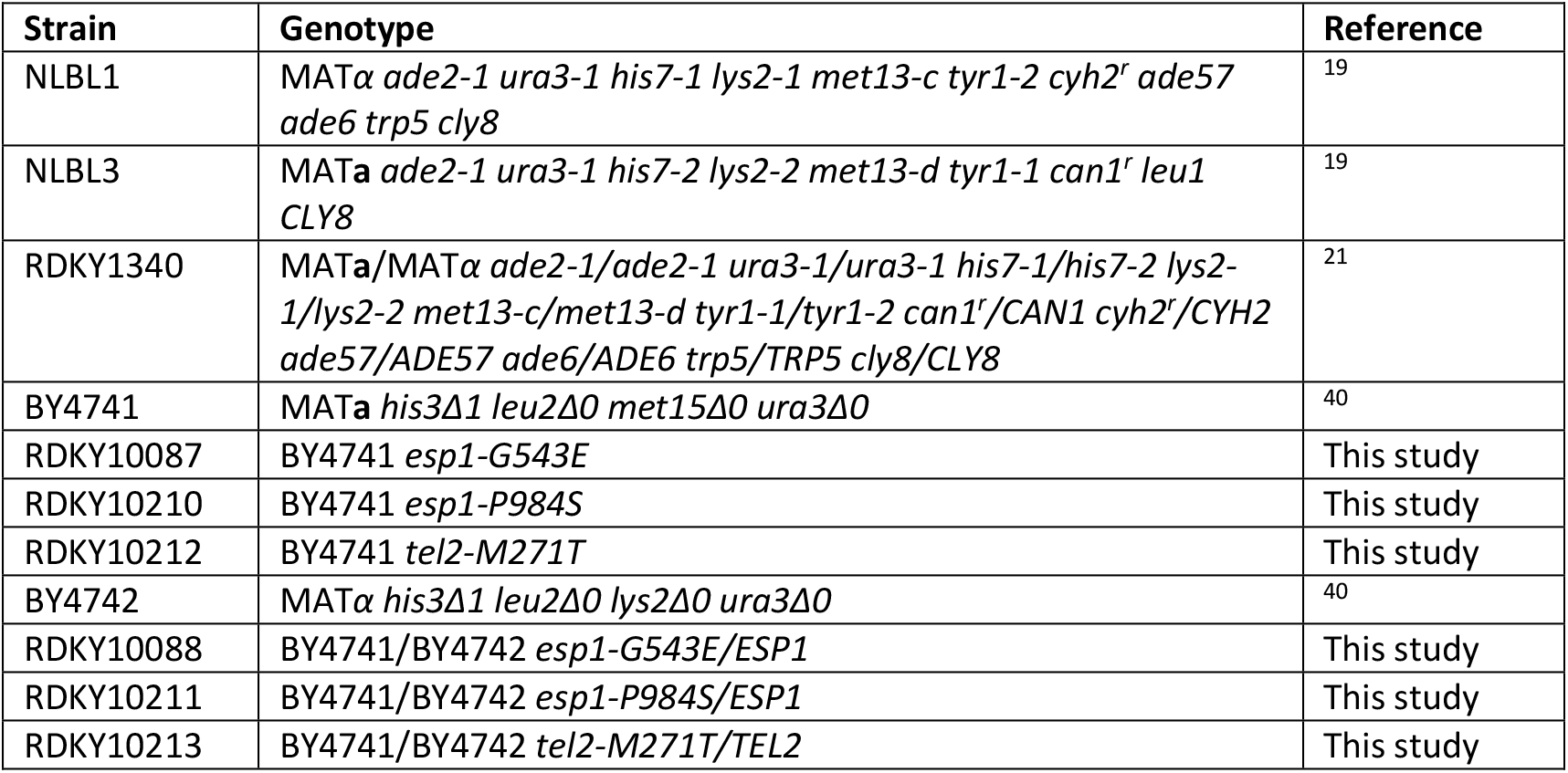
*S. cerevisiae* strains.

**Figure 1.**
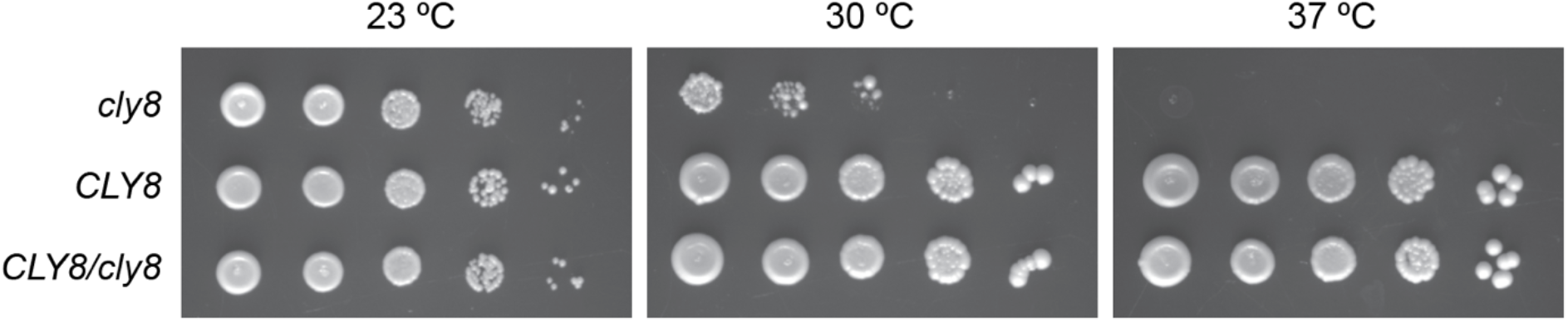
The *cly8* mutation causes a recessive temperature-sensitive growth phenotype. The *cly8* (NLBL1) and *CLY8* (NLBL3) haploid strains and the *cly8/CLY8* (RDKY1340) diploid strain were grown overnight at 23°C, subjected to 10-fold serial dilutions, and spotted onto YPD plates. Plates were grown at 23°C, 30°C, or 37°C as indicated. Images are representative plates from 3 experiments.

### Whole genome sequencing of *cly8* strains identifies several candidate missense mutations

Genetic mapping localizes *cly8* to the right arm of chromosome VII in the ∼80 kbp between *GCD2* and *SPT6* **(Fig. 2A)** ^8,12-16^; however, none of the genes in this interval have a direct role in maintaining the cell wall, reminiscent of the previously identified *cly* genes *GPI2, PKC1*, and *PSA1*. To identify candidate *cly8* sequence variants, the *cly8* haploid strain NLBL1, the *CLY8* haploid strain NLBL3, and the *CLY8/cly8* heterozygous diploid strain RDKY1340 were analyzed by paired-end whole genome sequencing. The average sequencing depths for the strains were 99, 110, and 104 reads, respectively **(Table 2)**. Using this data, we were able to identify the known mutations in these strains **(Table 3)**.

**Table 2.**
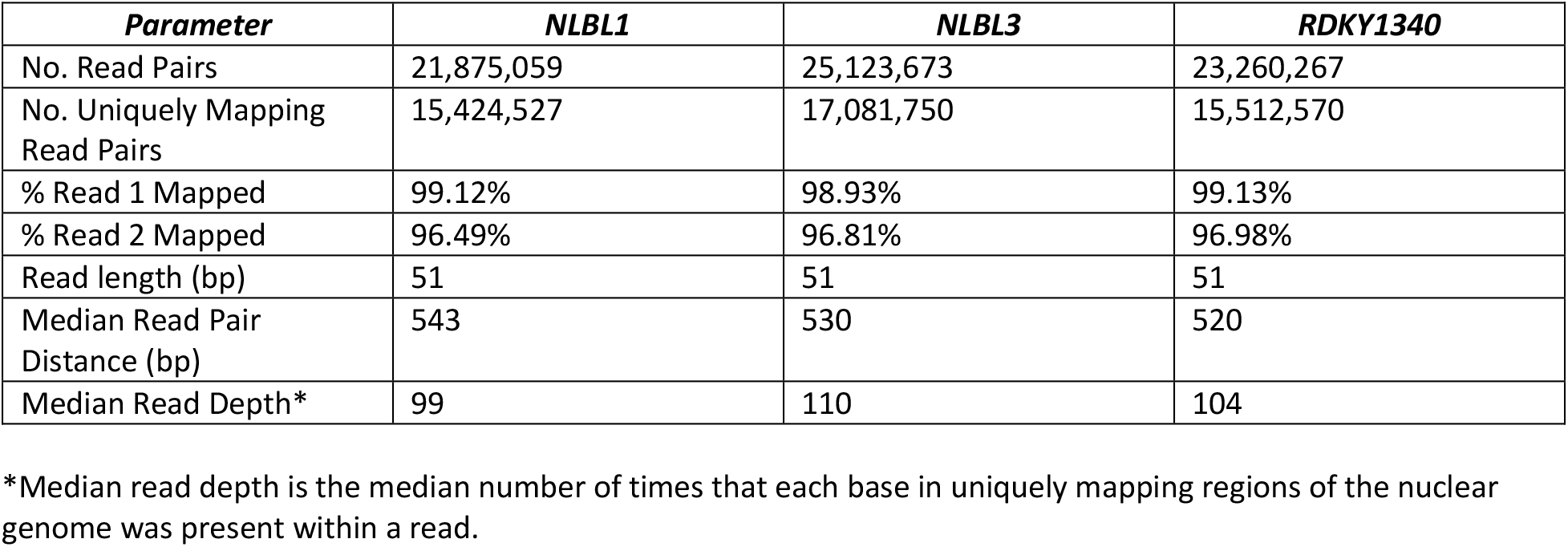
Whole genome sequencing statistics.

| ***Parameter*** | ***NLBL1*** | ***NLBL3*** | ***RDKY1340*** |
| --- | --- | --- | --- |
| No. Read Pairs | 21,875,059 | 25,123,673 | 23,260,267 |
| No. Uniquely Mapping Read Pairs | 15,424,527 | 17,081,750 | 15,512,570 |
| % Read 1 Mapped | 99.12% | 98.93% | 99.13% |
| % Read 2 Mapped | 96.49% | 96.81% | 96.98% |
| Read length (bp) | 51 | 51 | 51 |
| Median Read Pair Distance (bp) | 543 | 530 | 520 |
| Median Read Depth* | 99 | 110 | 104 |

\*Median read depth is the median number of times that each base in uniquely mapping regions of the nuclear genome was present within a read.

**Table 3.** Identification of reported mutations in strains NLBL1, NLBL3, and RDKY1340.

| ***Chromosomal location*** | ***Mutation*** | ***NLBL1*** | ***NLBL3*** | ***RDKY1340*** |
| --- | --- | --- | --- | --- |
| chrII:470,200 | *lys2-2* | - | c.3727C>T<br>Q1243X | c.3727C/c.3727C>T<br>Q1243/Q1243X |
| chrII:470,320 | *lys2-1* | c.3607C>T<br>Q1203X | - | c.3607C>T/c.3607C<br>Q1203X/Q1203 |
| chrII:570,476 | *tyr1-1* | - | c.725T>A<br>L242X | c.725T/c.725T>A<br>L242/L242X |
| chrII:570,795 | *tyr1-2* | c.406A>T<br>K136X | - | c.406A>T/c.406A<br>K136X/K136 |
| chrII:715,994 | *his7-2* | - | c.479delA | c.472A/c.479delA<br>K158/frameshift |
| chrII:716,188 | *his7-1* | c.278C>T<br>S93F | - | c.278C>T/c.278<br>S93F/S93 |
| chrV:33,328 | *can1<sup>r</sup>* | - | c.139A>T<br>K47X | c.139A/c.139A>T<br>K47/K47X |
| chrV:116,867 | *ura3-1* | c.701G>A<br>G234E | c.701G>A<br>G234E | c.701G>A/c.701G>A<br>G234E/G234E |
| chrVII:56,707 | *ade57* | c.226G>C<br>G76R | - | c.226G>C/c.226G<br>G76R/G67 |
| chrVII:272,815 | *met13-d* | - | c.296T>A<br>L99X | c.296T/c.296T>A<br>L99/L99X |
| chrVII:273,733 | *met13-c* | c.1214T>A<br>L405X | - | c.1214T>A/c.1214T<br>L405X/L405 |
| chrVII:311,589 | *cyh2<sup>r</sup>* | c.112C>G<br>Q38E | - | c.112C>G/c.112C<br>Q38E/Q38 |
| chrVII:448,499 | *trp5* | c.37A>T<br>K13X | - | c.37A>T/c.37A<br>K13X/K13 |
| chrVII:478,561 | *leu1* | - | c.92insT | c.92T/c.92insT<br>L31/frameshift |
| chrVII:614,898 | *ade6* | c.1068C>A<br>D356E | - | c.1068C>A/c.1068C<br>D356E/D356 |
| chrXV:566,002 | *ade2-1* | c.190G>T<br>E64X | c.190G>T<br>E64X | c.190G>T/c.190G>T<br>E64X/E64X |

**Figure 2.**
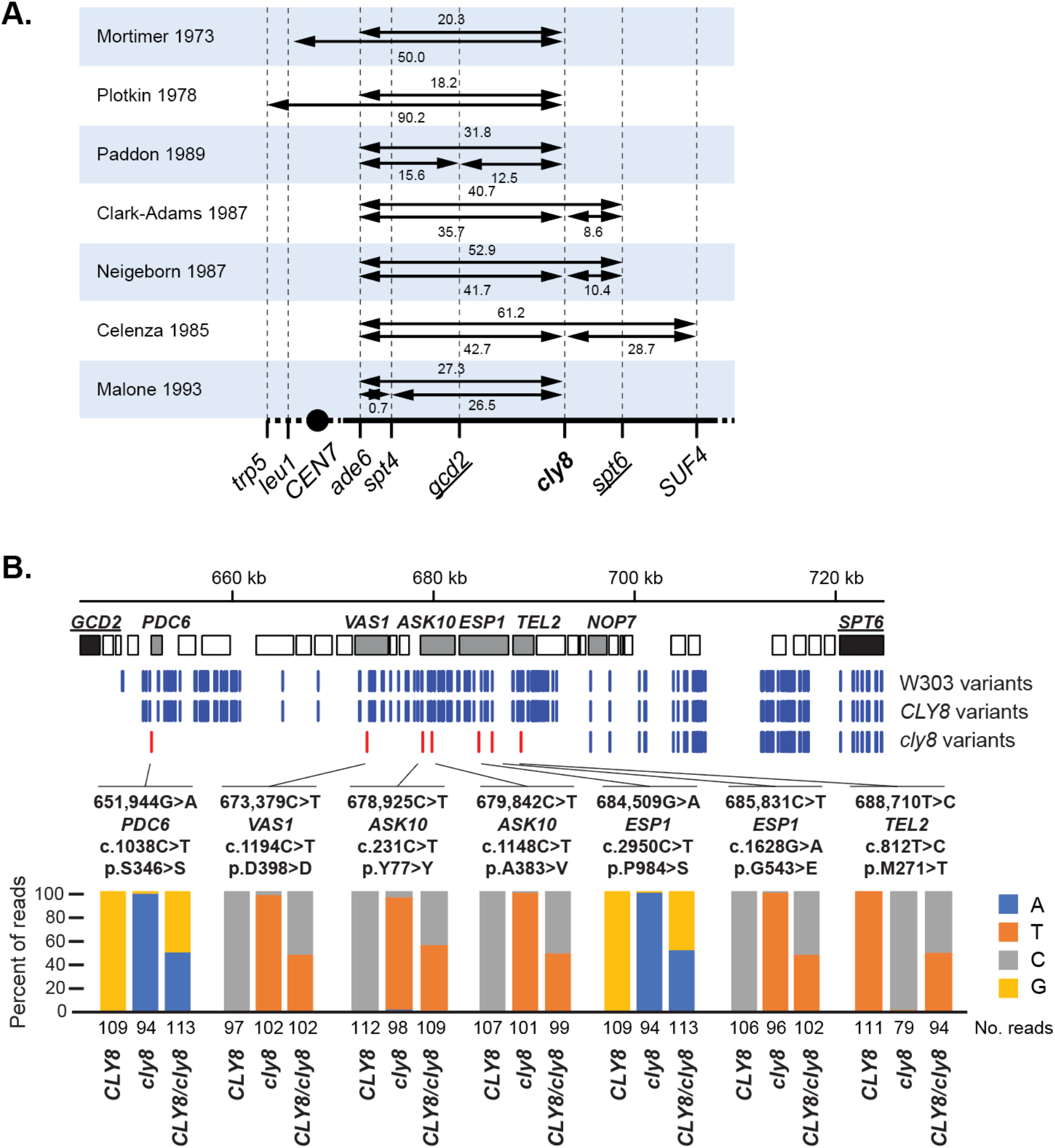
Identification of candidate *cly8* variants. **A**. Summary of previous genetic mapping of *cly8* indicates that it is between *gcd2* and *spt6*. Distances shown are in centiMorgans. **B**. Diagram of the position of variants identified by whole genome sequencing in the *GCD2-SPT6* region for the *cly8* and *CLY8* haploid strains. (Top) Diagram of all genes between *GCD2* and *SPT6* (black boxes). Blue vertical lines indicate the position of sequence variants in the W303 genome relative to the S288c reference genome. Red vertical lines indicate sequence variants only present in the *cly8* haploid strain. (Bottom) For each *cly8*-specific variant, the percent of reads called for each base and the effect of each variant is shown.

Sequence variants in the *GCD2-SPT6* region were analyzed in all three strains. The original *cly* mutants were isolated through mutagenesis of the A364A strain, which is closely related to the S288c reference strain ^1^. Remarkably, both the *cly8* and *CLY8* haploids shared chromosomal blocks sharing many single nucleotide variants (SNVs) with the *S. cerevisiae* W303 strain (**Fig. 2B**), which has approximately 9,500 SNVs relative to S288c ^22^. This suggests that the haploid strains were constructed through a cross to a W303-derived strain and that the *cly8* sequence variant (or variants) are likely present in the ∼44 kbp *PDC6*-*NOP7* region, as this region was derived from the A364A background in the *cly8* strain and from the W303 background in the *CLY8* haploid strain (**Fig. 2B**).

Candidate *cly8* variants in the ∼80 kbp *GCD2-SPT6* region were identified using the criteria that: [1] the variant must be a sequence change in the *cly8* strain relative to the S288c reference genome and the A364A genome, [2] the variant must not be present in the *CLY8* strain, and [3] the variant must be heterozygous in the *cly8/CLY8* diploid strain. These criteria were intended to filter out variants in these strains unrelated to the *cly8* mutation and identified 7 SNVs in the ∼44 kbp *PDC6*-*NOP7* region: 3 silent mutations in *PDC6, VAS1*, and *ASK10*, 1 missense mutations in the non-essential gene *ASK10* (A383V), and 3 missense mutations in the essential genes *ESP1* (G543E and P984S), and *TEL2* (M271T) **(Fig. 2B)**. All candidate *cly8* SNVs were GC>AT transitions, consistent with the preference of the N-methyl-N’-nitro-N-nitrosoguanidine alkylating agent used to generate the *cly8* mutation ^1,23^. Since *cly8* causes inviability at the non-permissive temperature, we focused on the missense mutations in essential genes *ESP1*, which encodes separase that cleaves the mitotic cohesin subunit Mcd1 to allow for chromosome segregation during mitosis ^24,25^, and *TEL2*, which encodes an ASTRA complex component that acts in telomere length regulation, chromatin remodeling, and folding of protein kinases ^26-28^.

### The *esp1-G543E* mutation confers temperature sensitive growth and chromosome segregation defects

The *esp1-G543E, esp1-P984S*, and *tel2-M271T* mutations were introduced individually into the wild-type strain BY4741 using homology-directed repair of CRISPR/Cas9-induced double stranded DNA breaks and were verified by PCR amplification and Sanger sequencing. The haploid *esp1-G543E* strain had substantial growth defects at both 30°C and 37°C (**Fig. 3A**) like the *cly8* strain **(Fig. 1**), whereas the *esp1-G543E/ESP1* diploid strain had normal growth at all temperatures tested (**Fig. 3A**). These results indicated that *esp1-G543E* mutation was recessive like *cly8*. In contrast, the haploid *esp1-P984S* and *tel2-M271T* strains did not have growth defects at elevated temperatures (**Fig. 3A**). The effects of these mutations were consistent with structural modeling: the Esp1-P984S and Tel2-M271T amino acid substitutions are predicted to be tolerated by the protein structure, whereas the Esp1-G543E is predicted to destabilize the Esp1 protein due to steric collisions with adjacent residues and potential burial of a charged residue in a hydrophobic space (**Fig. 4**).

**Figure 3.**
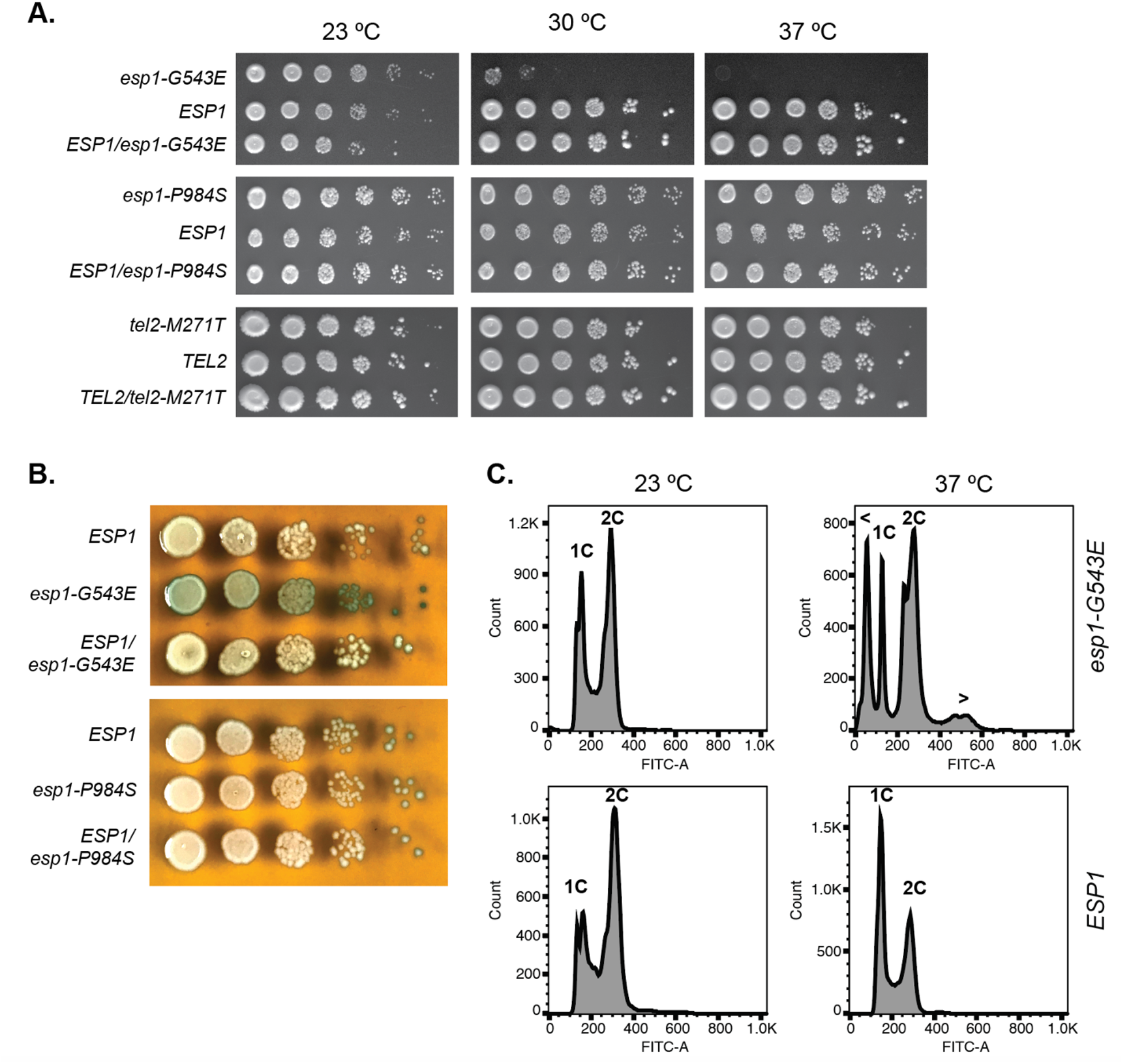
The *esp1-G543E* mutation causes temperature-sensitive defects in growth and chromosome segregation. **A**. 10-fold serial dilutions of the wildtype strain, the *esp1-G543E, esp1-P984S*, or *tel2-M271T* single mutant strains, and heterozygous mutant diploids strains spotted onto YPD plates and grown at 23°C, 30°C, or 37°C as indicated. Images are representative plates from 3 experiments. **B**. Colony overlay assay detecting increased cell lysis at 37°C for the *esp1-G543E* strain relative to the wildtype haploid strain or the *esp1-G543E/ESP1* diploid strain. The *esp1-P984S* mutation does not cause cell lysis. Cell lysis is detected through release of the vacuolar membrane protein alkaline phosphatase, which cleaves the chromogenic substrate BCIP and generates a blue color. Images are representative from 2 experiments. **C**. Histograms of the DNA content as measured by flow cytometry of BY4741 *esp1-G543E* and BY4741 haploid strains after 2.5 hours of growth at either 23°C or 37°C as indicated. “1C” = cells with 1C DNA content, “2C” = cells with 2C DNA content, “<“ = cells with less than 1C DNA content, and “>“ = cells with greater than 2C DNA content.

**Figure 4.**
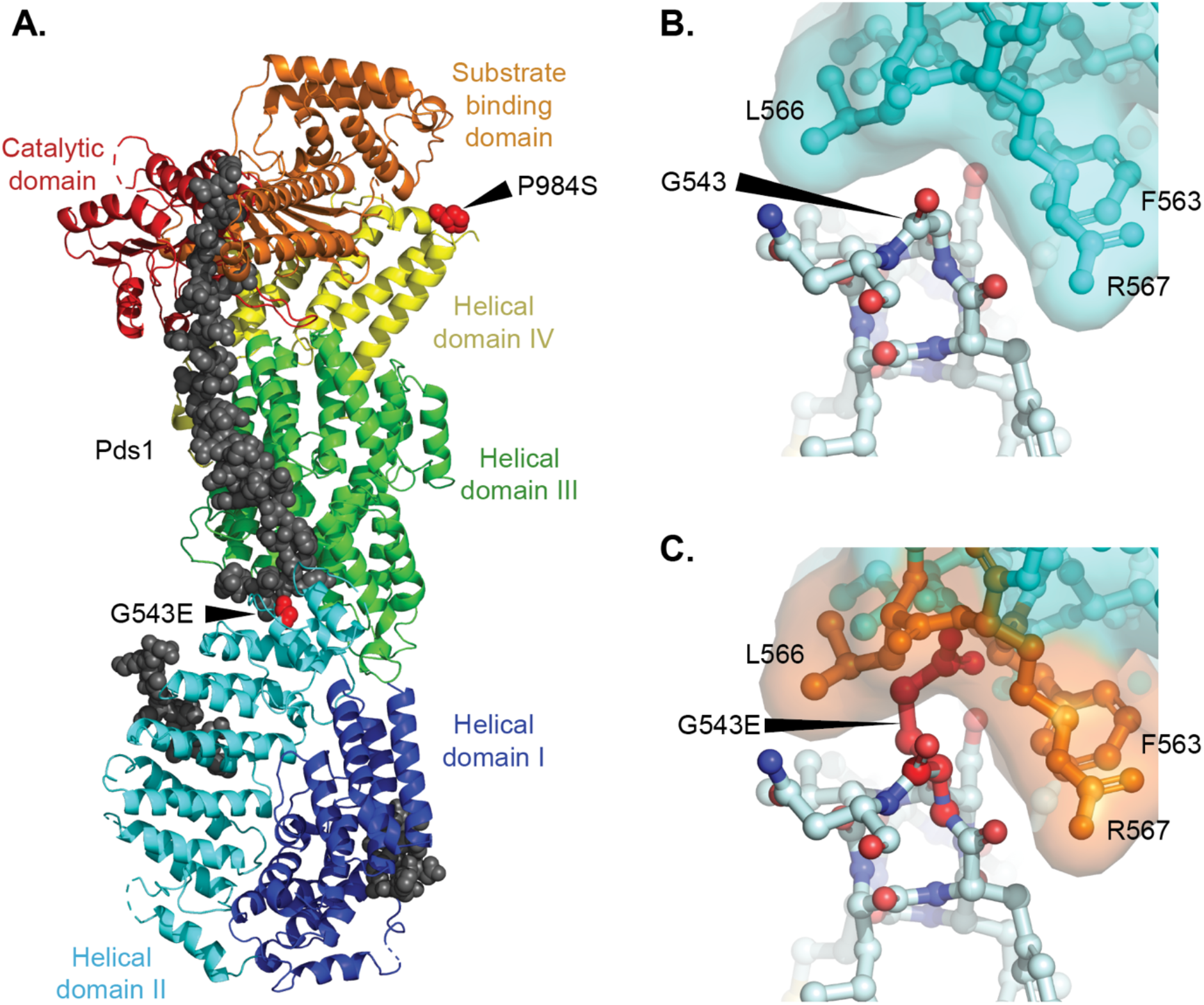
Structural modeling suggests that the Esp1-G543E causes steric collisions in helical domain II. **A**. Full length structure of the *S. cerevisiae* Esp1 (colored ribbon diagram) in complex with its Pds1 inhibitor (grey spheres; PDB id 5u1s; ^38^). The positions of the G543 and P984 residues are shown as red spheres. **B**. Close up of the interface between the packed helices containing Esp1-G543; the molecular surface of the adjacent Esp1 helix is shown. **C**. Close up of the interface where the Esp1-G543E mutation is modeled. All possible G543E rotamers cause substantial steric collisions with the adjacent helix (the side chain penetrates the molecular surface of the helix) and potentially places a charged residue into a hydrophobic environment. In contrast, the Esp1-P984S is predicted to be tolerated in which P984, which is at the N-terminus of a helix is replaced with a serine. Similarly, the Tel2-M271T is also predicted to be tolerated in the *S. cerevisiae* Tel2 structure (PDB id 3o4Z; ^39^), as M271 is at the end of a helix within the irregular N-terminal helical solenoid in which replacement by threonine ought to be tolerated based on size and the local environment of the side chain.

We then tested the *esp1-G543E* haploid strain for increased cell lysis at 37°C using a colony overlay assay where release of the vacuolar alkaline phosphatase converts a chromogenic substrate to a colored product. We found that the *esp1-G543E* strain had increased lysis at 37°C relative to the wildtype haploid strain and the *esp1-G543E/ESP1* diploid strain (**Fig. 3B**). In contrast and consistent with a lack of temperature sensitivity, the *esp1-P984S* strain did not show increased cell lysis relative to the wildtype haploid or the *esp1-P984S/ESP1* diploid strain (**Fig. 3B**).

The release of temperature-sensitive *esp1-1* cells synchronized with α-factor into medium at the non-permissive temperature causes an accumulation of cells with sub-1C and super-2C DNA content as revealed by flow cytometry analysis, consistent with defects in the cleavage of cohesin and proper chromosome segregation ^29^. To determine if the temperature sensitivity caused by the *esp1-G543E* mutation was also associated with a chromosome segregation defect, we transferred log-phase wildtype and *esp1-G543E* cells growing at 23°C to medium at either 23°C or 37°C for 2.5 h. Cells were then fixed and their DNA content was analyzed by flow cytometry (**Fig. 3C**). Both strains showed normal DNA content profiles after 2.5 h at 23°C with peaks at both 1C and 2C DNA content, which corresponded to roughly 22% and 50% of the cells, and roughly 22% of the cells with a DNA content between 1C and 2C. Shifting the cells to 37°C for 2.5 h caused the wildtype cells to accumulate with increased 1n DNA content (40% of the cells); however, the majority of cells maintained a normal DNA content. In contrast, when the *esp1-G543E* cells were shifted to 37°C, a substantial proportion of cells (>30%) had a DNA content with less than 1C or greater than 2C DNA content. These results demonstrate that *esp1-G543E* causes a temperature-sensitive chromosome segregation defect, which is reminiscent of the chromosome nondisjunction observed with the *esp1-1* mutation ^29^.

## DISCUSSION

Despite the potential utility of temperature-sensitive cell lysis mutations for the development of genetic assays ^19-21^, the identity of most of these mutations has not been established. Here, previous genetic mapping of the *cly8* mutation ^3,8,12-18^ was combined with whole genome sequencing to identify three candidate *cly8* sequence variants in two essential genes, *ESP1* and *TEL2*. Only the *esp1-G543E* mutation recapitulated the recessive temperature-sensitive growth profile of the starting *cly8* mutant strain when re-engineered into a wildtype strain, and the mutation also caused a temperature-sensitive cell lysis phenotype and a temperature-sensitive chromosome segregation defect. Taken together, these results indicate that the *cly8* mutation is an allele of *ESP1* and that introduction of the *esp1-G543E* mutation is sufficient to engineer the *cly8* growth defect for use in genetic assays.

The fact that *cly8* is an allele of *ESP1* might also explain unexpected genetic mapping results for crosses of *cly8* with *esp1-1* that gave a distance of 54.6 centiMorgans ^8^, which indicates independent segregation and is inconsistent with the genetic mapping of *cly8* and *esp1-1* with all of the other adjacent markers on chromosome VII (**Fig. 2A** and ^8^). Two alleles of the same gene normally would be expected to have a genetic distance close to 0 centiMorgans (e.g. the mutations never co-segregate to the same haploid progeny as they are allelic), but normal segregation in the *cly8/esp1-1* compound heterozygote is likely complicated by the lack of a wildtype copy of *ESP1*, which could lead to abnormal chromosome segregation.

The identification of an *esp1* mutation as giving rise to a cell lysis defect is surprising, given that Esp1 cleaves the cohesin subunit Mcd1 whereas the previously identified *cly* genes are more directly involved in cell wall biosynthesis and maintenance. Defects affecting the cohesin genes *IRR1, MCD1, SMC1*, and *SMC3*, which play crucial roles in chromosome segregation and the regulation of gene transcription, have been previously linked to cell wall defects, such as sensitivity to the chitin-binding compound calcoflour white ^30,31^. In the case of *mcd1-1*, the cell wall defect appears to be caused by reduced expression of cell wall biosynthesis and maintenance genes ^30^. Unlike *mcd1-1, esp1-G543E* does not cause sensitivity to calcofluor white (**Fig. 5**) nor is *ESP1* linked to changes in gene expression.

**Figure 5.**
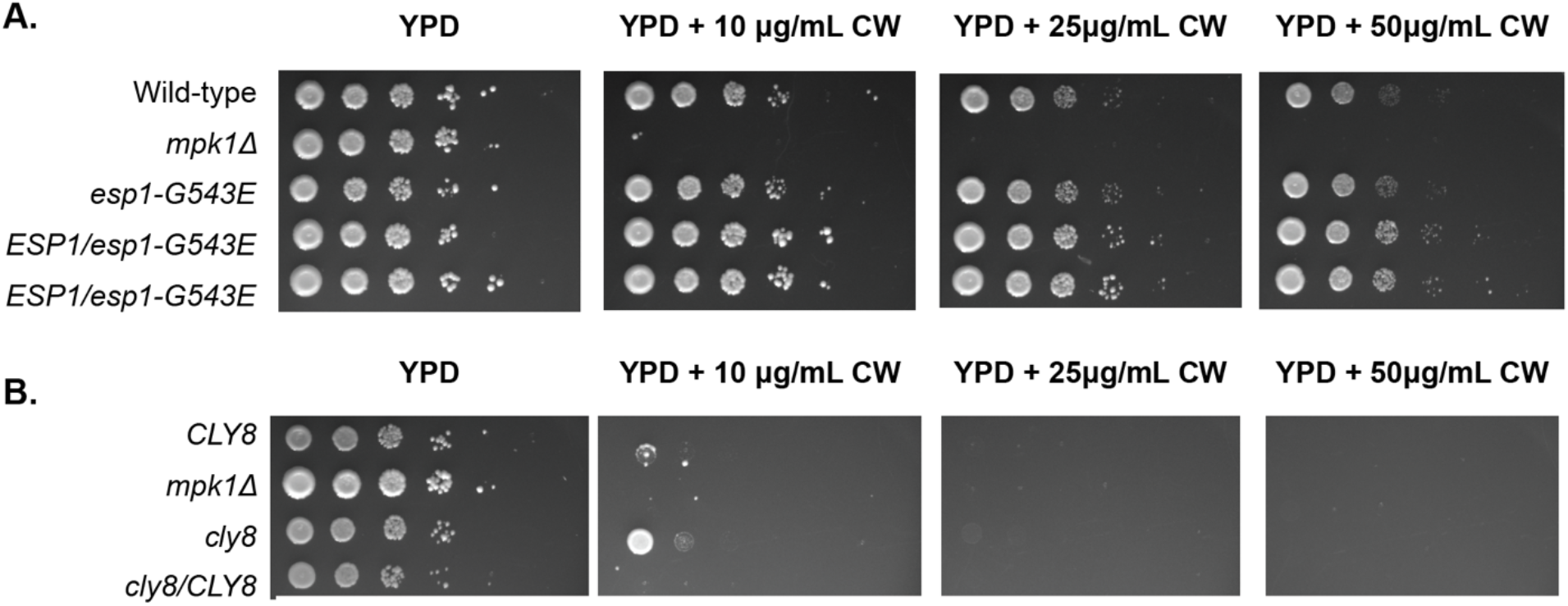
The *esp1-G543E* mutation does not cause sensitivity to cell wall stress caused by calcofluor white (CW). **A**. 10-fold serial dilutions of BY4741, BY4741 *mpk1Δ*, BY4741 *esp1-G543E*, and the BY4741/BY4742 *ESP1/esp1-G543E* diploid were plated on YPD and YPD containing 10, 25, and 50 μg/mL CW. The *mpk1Δ* strain was CW-sensitive as expected, whereas the *esp1-G543E* mutant was not. **B**. 10-fold serial dilutions of the *CLY8* haploid strain NLBL3, the *mpk1Δ* strain, the *cly8* haploid strain NLBL1, and the *CLY8/cly8* diploid strain RDKY1340 were plated on YPD and YPD containing 10, 25, and 50 μg/mL CW. Like the *mpk1Δ* strain, NLBL1, NLBL3, and RDKY1340 were sensitive to CW even at 10 μg/mL, regardless of the *cly8* status. Together, these results indicates that CW sensitivity in these strains is caused by a mutation other than *cly8*.

Remarkably, deletion of the *FAB1* gene, which encodes a vacuolar membrane 1-phosphatidylinositol-3-phosphate 5-kinase, causes many defects, including a growth defect at 37°C and a cell lysis phenotype ^32^. Moreover, *fab1* mutations (<u>F</u>orms <u>A</u>ploid and <u>B</u>inucleate cells) leads to the accumulation of cells with less than 1C DNA content and greater than 2C DNA content due to a misalignment of the mitotic spindle caused by increased vacuole size ^32^. These phenotypes are strongly reminiscent of the chromosome segregation defects caused by *esp1* mutations and the temperature-sensitive cell-lysis defect caused by *cly8*, suggesting that cell lysis in *cly8* and *fab1* mutants occurs secondarily to chromosome mis-segregation. Intriguingly, the cell lysis *cly3* mutation has been mapped to a genomic region containing *FAB1* (chromosome VI between *SUP11* and *HIS2*), raising the possibility that that *cly3* could be a mutation of *FAB1* (**Fig. 6**; ^33^). The identification of the *cly8* mutation here allows for its use in genetic assays and suggests that the cell lysis phenotype is not only caused directly by cell wall maintenance and biosynthesis defects but also by some kinds of chromosome segregation defects.

**Figure 6.**
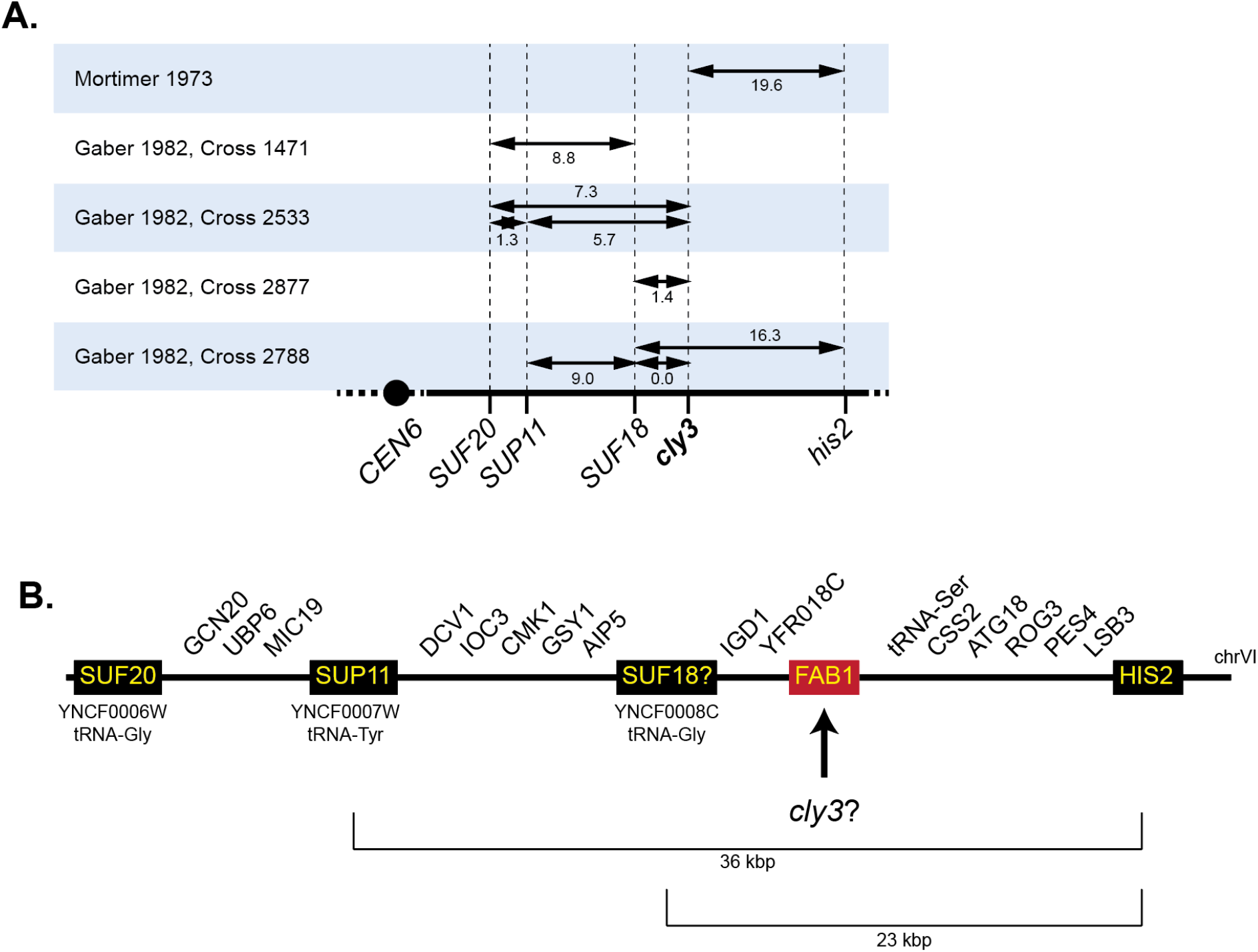
The *cly3* mutation may be a mutation of *FAB1*. **A**. Fine genetic mapping of *cly3* on the right arm of chromosome VI suggests that it lies in the 36 kbp region between *SUP11* and *HIS2* ^3,33^. **B**. Genes present in the *SUF11-HIS2* region of chromosome VI are the candidate sites for the *cly3* mutation. The *cly3* mutation is tightly linked to *SUF18*, which is a glycine tRNA mutation that suppresses +G frameshift mutation in some glycine codons, such as 5’-GGGU-3’ in the *his4-38* allele ^33^. *SUF18* has not been mapped, but a glycine tRNA (*YNCF0008C*, previously called tG(GCC)F2) between *AIP5* and *IGD1* is the only candidate in the *SUP11-HIS2* region in the *S. cerevisiae* reference genome. This identification of *SUF18* would further restrict *cly3* to the left end of the 23 kbp *YNCF008C-HIS2* region. None of the genes in this region are essential, and the only gene reported to cause a temperature-sensitive defect when deleted is *FAB1*, which has pleotropic effects due to vacuolar dysfunction, including sharing a cell lysis phenotype and a chromosome segregation defect phenotype with *cly8* ^32^. Notably, *FAB1* is much closer to *YNCF0008C* than *HIS2*, as would be predicted from the *cly3* genetic mapping results.

## METHODS

### Plasmid construction

The plasmids encoding Cas9 and a single guide RNA (sgRNA) to target cleavage adjacent to *ESP1*-G543, *ESP1*-P984, and *TEL2*-M271 were generated by annealing oligonucleotides encoding the guide sequence and ligating them using T4 DNA ligase (New England Biolabs) into SapI digested pRS425-Cas9-2xSapI. The pRS425-Cas9-2XSapI vector was constructed in Bruce Futcher’s laboratory (State University of New York, Stoney Brook). Introduction of the gRNA targeting sequence was verified by Sanger sequencing (Retrogen). The pRDK2100 plasmid for cutting next to *ESP1*-G543 was constructed using the oligonucleotides 5’-atcATTATGCTTTCAGTACTGTA-3’ and 5’-aacTACAGTACTGAAAGCATAAT-3’ (uppercase letters correspond to the guide sequence and complement). The pRDK2141 plasmid for cutting next to *ESP1*-P984 was constructed using the oligonucleotides 5’-atcTCAAACATACCTTTGAAGAA-3’ and 5’-aacTTCTTCAAAGGTATGTTTGA-3’. The pRDK2142 plasmid for cutting next to *TEL2*-M271 was constructed using the oligonucleotides 5’-atcTAACAAACTTCGTGGAAGAG-3’ and 5’-aacCTCTTCCACGAAGTTTGTTA-3’.

### Strain construction

Haploid single mutant strains were created by transforming the wild-type strain BY4741 with the plasmid encoding Cas9 and the sgRNA with a homologous recombination target generated by annealing two oligonucleotides. These homologous recombination targets introduced the mutation as generated a silent mutation disrupting the PAM site. Leu^+^ transformants were isolated and the mutated region was amplified by PCR and subjected to Sanger sequencing (Retrogen). The *esp1-G543E* strain RDKY10087 was created with pRDK2100 and a homologous recombination target generated by annealing the oligonucleotides 5’-CGA GTT TTC AAA CAG TTC AGA AAT TAT GCT TTC AGT ACT GTA TGa AAA TTC ATC TAT CGA AAA CAT TCC TTC CGA AAA TTG GTC-3’ and 5’-GAC CAA TTT TCG GAA GGA ATG TTT TCG ATA GAT GAA TTT tCA TAC AGT ACT GAA AGC ATA ATT TCT GAA CTG TTT GAA AAC TCG-3’ (lower case base corresponds to the sequence change that introduced the mutation and disrupted the PAM sequence). The *esp1-P984S* strain RDKY10210 was created with pRDK2141 and a homologous recombination target generated by annealing the oligonucleotides 5’-ACA CAA TAT GTG GCA AAA AGT AAT GAG CCA ATT GGA GGA AGA TtC TTT CTT CAA AGG TAT GTT TGA ATC TAC CCT TGG GAT TCCC-3’ and 5’-GGG AAT CCC AAG GGT AGA TTC AAA CAT ACC TTT GAA GAA AGa ATC TTC CTC CAA TTG GCT CAT TAC TTT TTG CCA CAT ATT GTGT-3’ (lower case base corresponds to the sequence change that introduced the mutation and disrupted the PAM sequence). The *tel2-M271T* strain RDKY10212 was created with pRDK2142 and a homologous recombination target generated by annealing the oligonucleotides 5’-GAA ATA CAG TCG CTG CCA TTG AAG GAA GTG ATA GTA CGT CTG AcG AGC AAt CAC TCT TCC ACG AAG TTT GTT AGC GCT TTGG-3’ and 5’-CCA AAG CGC TAA CAA ACT TCG TGG AAG AGT GaT TGC TCg TCA GAC GTA CTA TCA CTT CCT TCA ATG GCA GCG ACT GTA TTTC-3’ (lower case bases correspond to the sequence changes that introduced the mutation and disrupted the PAM sequence). Diploid strains were generating by crossing single mutant strains to the wild-type strain BY4742 (MAT*α his3Δ1 leu2Δ0 lys2Δ0 ura3Δ0*) to generate RDKY10088 (*esp1-G543E/ESP1*), RDKY10211 (*esp1-P984S/ESP1*), and RDKY10213 (*tel2-M271T/TEL2*).

### Whole genome paired-end sequencing

Multiplexed paired-end libraries were constructed from 2 μg of genomic DNA purified from NLBL1, NLBL3, and RDKY1340 using the Purgene kit (Qiagen). The genomic DNA was sheared using M220 focused-ultrasonicator (Covaris) and end-repaired using the End-it DNA End-repair kit (Epicentre Technologies). Common adaptors from the Multiplexing Sample Preparation Oligo Kit (Illumina) were then ligated to the genomic DNA fragments, and the fragments were then subjected to 18 cycles of amplification using the Library Amplification Readymix (KAPA Biosystems). The amplified products were fractionated on an agarose gel to select 600 bp fragments, which were subsequently sequenced on an Illumina NovaSeq 6000 using the Illumina GAII sequencing procedure for paired-end short read sequencing. Reads from each read pair were mapped separately by bowtie version 2.2.1 ^34^ to a reference sequence that contained revision 64 of the *S. cerevisiae* S288c genome (http://www.yeastgenome.org). Mapped reads were analyzed using pyrus version 0.7 (http://www.sourceforge.net/p/pyrus-seq) ^35^. Sequence data is available from National Center for Biotechnology Information Sequence Read Archive (SRA) under accession number PRJNA750757. Sequencing data for the W303 (SRR1569900) ^36^ and A364A strains (SRR3481429) ^37^ were downloaded from the SRA and analyzed with pyrus.

### Colony overlay assay

The colony overlay assay was performed essentially as described by Paravicini et al ^5^. Cells were spotted onto YPD plates and grown for 2 days at 30°C prior to shifting to 37°C for 18 hours. Plates were then overlaid with an alkaline phosphatase assay solution containing 0.05 M glycine hydrochloride pH 9.5, 1% agar, and 10 mM 5-bromo-4-chloro-3-indolyl-phosphate (BCIP) and photographed after 30 minutes. Colonies with lysed cells turn blue due to the release of the vacuolar membrane alkaline phosphatase.

### DNA content analysis

Strains to be analyzed were grown overnight in YPD at 23°C, and 0.5 mL were added to 5 mL of YPD medium followed by 1 h growth at 23°C. The cultures were then split in half, pelleted by centrifugation, and then resuspended into 5 mL of YPD pre-equilibrated at 23°C or 37°C and then grown at 23°C or 37°C for 2.5 hrs. Cells were then collected by centrifugation, washed in 50 mM sodium citrate (pH 7.5), and fixed by resuspending in 70% ethanol for 1 h. Fixed cells were then washed in 50 mM sodium citrate (pH 7.5), resuspended in the same buffer containing 0.25 mg/ml of RNaseA and 1 mg/mL proteinase K, and incubated at 37°C overnight. Cells were then pelleted and resuspended in 50 mM sodium citrate (pH 7.5) buffer containing 1 μM Sytox-green (Invitrogen) and incubated for at least 1 h prior to analysis. Samples were then analyzed using a Becton Dickinson Fortessa instrument using FACSDiva Software. DNA content was then analyzed using FlowJo version 10.

## ACKNOWLEDGEMENTS

The authors thank Michael Esposito and Richard Kolodner for gifts of strains. Illumina sequencing was conducted at the IGM Genomics Center, University of California, San Diego, La Jolla, CA utilizing an Illumina NovaSeq 6000 that was purchased with funding from a National Institutes of Health SIG grant (#S10 OD026929). This research was funded by the grant NIH R01 GM026017.

